# SARS-CoV-2 Vaccine Booster Elicits Robust Prolonged Maternal Antibody Responses and Passive Transfer Via The Placenta And Breastmilk

**DOI:** 10.1101/2022.11.29.518385

**Authors:** Nicole E. Marshall, Madison B. Blanton, Brianna M. Doratt, Delphine C. Malherbe, Monica Rincon, Heather True, Taylor Mcdonald, Caroline Beauregard, Reuben Adatorwovor, Ilhem Messaoudi

**Author notes:** Corresponding authors: Nicole Marshall and Ilhem Messaoudi, Nicole Marshall, Department of Obstetrics and Gynecology, Oregon Health & Science University, 3181 SW Sam Jackson Park Rd, Portland, OR 97239, Ilhem Messaoudi, Department of Microbiology, Immunology, and Molecular Genetics, College of Medicine, University of Kentucky, 760 Press Ave, Lexington, KY 40536.

## Abstract

**Background:** Infection during pregnancy can result in adverse outcomes for both pregnant persons and offspring. Maternal vaccination is an effective mechanism to protect both mother and neonate into post-partum. However, our understanding of passive transfer of antibodies elicited by maternal SARS-CoV-2 mRNA vaccination during pregnancy remains incomplete.

**Objective:** We aimed to evaluate the antibody responses engendered by maternal SARS-CoV-2 vaccination following initial and booster doses in maternal circulation and breastmilk to better understand passive immunization of the newborn.

**Study Design:** We collected longitudinal blood samples from 121 pregnant women who received SARS-CoV-2 mRNA vaccines spanning from early gestation to delivery followed by collection of blood samples and breastmilk between delivery and 12 months post-partum. During the study, 70% of the participants also received a booster post-partum. Paired maternal plasma, breastmilk, umbilical cord plasma, and newborn plasma samples were tested via enzyme-linked immunosorbent assays (ELISA) to evaluate SARS-CoV-2 specific IgG antibody levels.

**Results:** Vaccine-elicited maternal antibodies were detected in both cord blood and newborn blood, albeit at lower levels than maternal circulation, demonstrating transplacental passive immunization. Booster vaccination significantly increased spike specific IgG antibody titers in maternal plasma and breastmilk. Finally, SARS-CoV-2 specific IgG antibodies in newborn blood correlated negatively with days post initial maternal vaccine dose.

**Conclusion:** Vaccine-induced maternal SARS-CoV-2 antibodies were passively transferred to the offspring *in utero* via the placenta and after birth via breastfeeding. Maternal booster vaccination, regardless of gestational age at maternal vaccination, significantly increased antibody levels in breastmilk and maternal plasma, indicating the importance of this additional dose to maximize passive protection against SARS-CoV-2 infection for neonates and infants until vaccination eligibility.

## INTRODUCTION

The fetal immune system is highly immature resulting in heightened susceptibility to infection.^1-3^ Similarly, infection during pregnancy can lead to significant adverse outcomes for both pregnant persons and offspring^4^, as has been demonstrated by the SARS-CoV-2 global pandemic. These adverse outcomes can be mitigated through maternal vaccination which protects the pregnant person and the neonate/infant via passive transfer of maternal antibodies either *in utero* via the placenta or after birth via breastmilk.^5-9^ Immunoglobulins G (IgG) pass from maternal to fetal circulation via neonatal plasma Fc receptors (FcRN) in the placenta and fetal intestines.^9^

Pregnant persons are encouraged to receive the seasonal influenza vaccine as soon as it becomes available, regardless of gestational trimester,^8, 10^ to prevent maternal influenza infection. Babies born to mothers who were vaccinated against influenza during pregnancy have higher hemagglutination-inhibition antibody (HIA) titers.^11^ Similarly, influenza-specific antibody titers in breastmilk are higher in mothers who were vaccinated.^12^ Current recommendations also include administration of the tetanus toxoid, reduced diphtheria toxoid, and acellular pertussis (Tdap) vaccine at approximately 27-36 weeks’ gestation^13^ as prior studies of maternal vaccination have suggested that IgG is preferentially transported across the placenta in late gestation, resulting in neonatal levels higher than maternal plasma levels.^14^

The Centers of Disease Control and Prevention (CDC) now recommends vaccination against SARS-CoV-2 for persons who are pregnant or plan to become pregnant.^15^ Despite mounting evidence that maternal vaccination is safe, decreases maternal and neonatal morbidity and mortality, and leads to passive newborn immunization via both placental transfer and breastfeeding,^16-20^ there remains a high level of vaccine hesitancy,^21^ resulting in only 71.5% of the pregnant population receiving a SARS-CoV-2 vaccination^22^ and less than half receiving a booster dose.^23^ Additionally, 46% of pregnant women recorded vaccine hesitancy ^24^ citing safety concerns ^25^ despite lack of significant adverse gestational outcomes,^26^ a comparable antibody response in pregnant and nongravid females,^19^ evidence of transplacental passive transfer of IgG antibodies,^16^ and detectable antibody levels in breastmilk after the initial vaccination series.^20, 27^ Moreover, booster vaccinations led to increased levels of maternal IgG1 and IgA antibodies in umbilical cord blood^28^ and breastmilk.^29^

For some pregnant individuals, vaccination decisions are highly influenced by a primary goal to protect neonatal health. Thus, their decision as to whether to receive primary or booster vaccinations during pregnancy or to delay vaccination until a later gestational age or postpartum are shaped by knowledge about impact of vaccine timing and duration of protection. To date, there are limited published data to guide these decisions.^16, 18-20, 27-32^ Previous studies investigating maternal SARS-CoV-2 vaccination include minimal longitudinal sampling that spans across the initial vaccination series and booster. In addition, the impact of gestational age at the time of vaccination on maternal and fetal/newborn antibody titers remains poorly understood. In this study, we addressed these gaps in our knowledge by measuring antibody levels in maternal circulation, cord blood, newborn blood, and breastmilk, throughout gestation, at birth, and up to 12 months post-partum in a cohort of 121 women.

## MATERIALS AND METHODS

### Ethical statement

The study was approved by the institutional Ethics Review Boards of Oregon Health & Science University and the University of Kentucky. All subjects provided written consent prior to enrollment which occurred from March 2021 to June 2022.

### Sample processing

Breastmilk was diluted 1:1 in 1X HBSS (CORNING, Corning, NY) and centrifuged at 810g at room temperature for 10 minutes. After the removal of the fat layer, the supernatant was collected and stored at −80°C until analysis. Whole blood samples were processed as previously described.^33^

### Enzyme-linked immunosorbent assay (ELISA)

An indirect ELISA was used to determine the IgG end-point titer (EPT) of antibodies against SARS-CoV-2 receptor-binding domain (RBD) of the spike protein as described in ^34^. Newborn plasma was initially diluted 1:50 in blocking buffer (BB) while maternal plasma and umbilical cord plasma were initially diluted 1:30. Breastmilk was loaded at a 4:1 dilution to BB. A three-fold dilution series was performed for all plasma samples while breastmilk samples were not diluted further. Plasma endpoint IgG titers (EPT) were calculated using log-log transformation of the linear portion of the curve, and 0.1 OD units as cut-off. For breastmilk, antibody levels were reported as optical density (OD) values.

To measure specific IgG isotypes titers, newborn plasma was diluted 1:50 for IgG1 and IgG3, and 1:10 for IgG2 and IgG4 in BB while maternal plasma and umbilical cord plasma were diluted 1:30 for IgG1 and IgG3 and 1:10 for IgG2 and IgG4 in BB followed by 6 three-fold dilutions. Breastmilk was loaded at a 4:1 ratio of sample to BB. Responses were visualized by adding HRP anti-human IgG1, IgG2, IgG3, or IgG4 (1:4,000 in BB) (SouthernBiotech, Birmingham, Alabama). Plates were read and analyzed as previously cited ^34^.

### Statistical analyses

We conducted all statistical analyses using Prism9 (Graphpad Prism, San Diego, California) and SAS version 9.4 (TS1M1, SAS institute, Cary, NC) statistical software. Data was tested for normality. Normally and not normally distributed data sets were analyzed by parametric and nonparametric tests, respectively. Two group comparisons were conducted via a paired T-test if samples were from the same participants and an unpaired T-test if not. Multiple group comparisons were tested by a paired one-way ANOVA with Dunn’s multiple comparison when all data were derived from the same subjects. An unpaired ANOVA was used for IgG isotype analysis when data on for all four isotypes were not derived from the same group of subjects. Additionally, the half-life of the antibody response following initial or booster dose of SARS CoV-2 vaccines was calculated using the standard exponential decay rate formula. The half-life was estimated using a probability integral transform. Pearson’s correlation analysis was used to establish pair-wise relationships. P-values and FDR ≤ 0.05 were considered statistically significant; 0.05-0.1 were denoted as trending.

## RESULTS

### Cohort Description

Maternal blood and breastmilk samples were obtained longitudinally from 121 SARS-CoV-2 vaccinated participants (Pfizer BN162b2 or Moderna mRNA-1273). The overwhelming majority (90.9%) of participants received the Pfizer BN162b2 vaccine; the remainder received the Moderna mRNA-1273 vaccine. Newborn blood, umbilical cord blood, and colostrum were collected at the time of delivery (**Figure 1A**). The characteristics of the cohort are described in **Table 1**. Participants received their first vaccine either pre-pregnancy (12.4%); first trimester (T1, 12.4%); second trimester (T2, 29.8%); third trimester (T3, 22.3%); or postpartum (23.1%). Nearly three-quarters (72.7%) of participants received a booster dose post-partum (**Table 1**). The average maternal age at initial vaccination and the average gestational age at delivery were not different between groups.

**Table 1.** Cohort Metadata. **Cohort characteristics.** Subjects are stratified by the trimester of initial maternal SARS-CoV-2 vaccination. Maternal age and gestational age at delivery are mean ± standard deviation. There is no significant difference among maternal age nor gestational age of delivery within the cohort when stratified by vaccination timepoint.

|  | Pre-pregnancy | T1 | T2 | T3 | Post-Partum |
| --- | --- | --- | --- | --- | --- |
| All (121) |  |  |  |  |  |
| N (%) | 15 (12.4%) | 15 (12.4%) | 36 (29.8%) | 27 (22.3%) | 28 (23.1%) |
| Maternal age (years) | 34.9 ± 3.7 | 34.5 ± 3.1 | 34.2 ± 4.1 | 34.8 ± 4.7 | 33.6 ± 4.6 |
| Gestational age at delivery (years) | 38.9 ± 0.9 | 39.2 ± 1.2 | 38.9 ± 1.3 | 39.0 ± 2.0 | 39.0 ± 1.2 |
| Fetal sex (% female) | 33% | 40% | 47% | 52% | 36% |
| Initial Vaccine Series |  |  |  |  |  |
| Pfizer | 13 | 13 | 34 | 26 | 24 |
| Moderna | 2 | 2 | 2 | 1 | 4 |
| Received booster | 14 | 12 | 27 | 23 | 15 |
| Days between initial vaccination and booster | 278 ± 27.8 | 298.5 ± 27.7 | 242.3 ± 32.0 | 257.5 ± 32.8 | 263.3 ± 32.4 |

**Figure 1:**
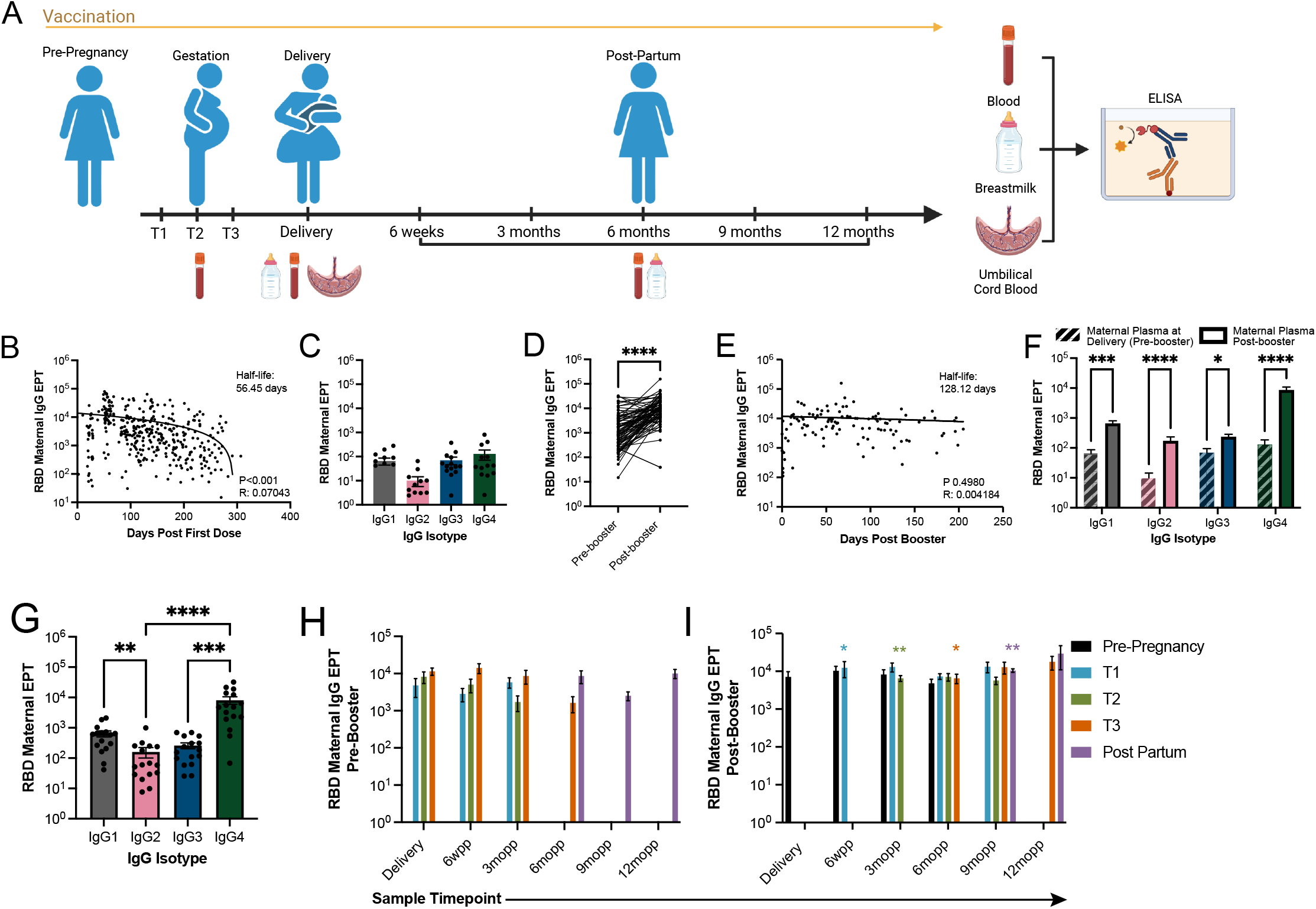
SARS-CoV-2 initial vaccination regimen and the booster result in robust RBD-specific IgG antibody response in maternal plasma. **(A)** Experimental design to investigate the impact of maternal SARS-CoV-2 vaccination on passive transmission of RBD-specific IgG antibodies by assessing antibody titers in maternal plasma, UCB, newborn plasma, and breastmilk. **(B)** RBD-specific IgG antibody titers in maternal plasma relative to days post first vaccination (n=370 samples). **(C)** IgG isotypes titers in maternal circulation 163.53 ± 14.28 days post first vaccine and pre-booster, (n=15). **(D)** RBD-specific IgG antibody titers 50.59 ± 4.46 days before and 55.74 ± 4.14 days after booster dose (n=77 pairs). **(E)** RBD-specific IgG antibody titers in maternal plasma relative to days post booster dose (n=112). **(F)** IgG isotype levels 84.07 ± 12.34 days before and 58.47 ± 8.98 days after the booster dose (n=15 pairs). **(G)** Isotype analysis of maternal plasma 58.13 ± 8.87 days post booster (n=16). **(H-I)** Average RBD-specific IgG antibody titers post-partum in women who have not received the booster **(H)** (Prepreg, n=0; T1, n=26; T2, n=57; T3, n=64; Postpartum, n=40) and **(I)** who have received the booster (Prepreg, n=30; T1, n=16; T2, n=43; T3, n=30; Postpartum, n=12) classified by trimester of initial vaccination. Bar graphs show median values with the standard error of the mean (SEM). * indicates a significant difference between the antibody levels at that timepoint when comparing pre- and post-booster groups **(panel H vs. panel I)**. ∗ *p* < 0.03 ∗∗ *p* < 0.002, ∗∗∗ *p* < 0.0002, ∗∗∗∗ *p*<0.0001.

### Vaccination against SARS-CoV-2 leads to a robust antibody response in pregnant women which is significantly increased following booster dosing

RBD-specific IgG titers strongly inversely correlated with time elapsed since the first vaccination (r=0.07043 *p*<0.0001) with a half-life of 56.45 days. (**Figure 1B)**. IgG effector function has been shown to vary among antibody isotypes.^35^ Specifically, viral infections are associated with increased IgG1 and IgG3 isotypes, while significant IgG2 involvement is linked to defense mechanisms against bacterial capsular polysaccharides.^35, 36^ IgG4 immunoglobulin responses have been attributed to subsequent or persistent exposure to antigen.^35,36,37^ Therefore, we investigate the IgG isotype specificity post mRNA vaccination. The initial series elicited a comparable response among all IgG isotypes (**Figure 1C)**. Circulating RBD-specific IgG titers increased significantly after the booster dose (*p*<0.0001) (**Figure 1D**). In addition, antibody responses elicited by the booster dose had a longer half-life of 128.12 days (**Figure 1B, 1E**) with minimal decrease in IgG levels after 6 months. The booster led to a significant increase in all four IgG isotypes measured in maternal plasma (**Figure 1F)** with IgG4 becoming the dominant isotype **(Figure 1G)**. Gestational age at the time of initial vaccination did not significantly impact antibody levels in maternal plasma at delivery and post partum (**Figure 1H)**. However, RBD-specific IgG antibody titers in women who were subsequently boosted were significantly higher 6 weeks post-partum (wpp) (*p*=0.0272), 3 months post-partum (mopp) (*p=*0.0032), 6mopp (0.0467), and 9mopp (0.0089) respectively relative to titers in women who were not boosted **(Figure 1I**).

### Breastmilk IgG antibody levels positively correlated with maternal circulating antibody levels

The initial 2-dose vaccination regimen resulted in detectable IgG antibody response in breastmilk (albeit much reduced levels compared to maternal plasma) with a half-life of 61.34 days (**Figure 2A)**. In contrast to maternal plasma, levels of RBD-specific IgG4 were significantly lower than those of IgG1 (p<0.0001) and IgG3 (*p* = 0.0011) in breastmilk after initial maternal vaccination (**Figure 2B)**. As described for maternal circulation, the booster led to a significant increase in breastmilk antibody levels (*p<*0.0001) (**Figure 2C**) and half-life (124.67 days) (**Figure 2D)**. After the booster dose, levels of IgG1 and IgG4 increased significantly (**Figure 2E**) with IgG4 becoming the dominant IgG isotype in breastmilk (**Figure 2F**). As seen for maternal systemic antibodies, gestational age at the time of initial vaccination did not significantly impact antibody levels in breastmilk (**Figure 2G**). However, booster vaccination resulted in a significant increase in RBD-specific IgG titers at 6wpp and 3mopp in women who were initially vaccinated in T1 (*p=*0.0055; *p=*0.056) or T2 (*p<*0.001 for both time points) compared to levels in women who had not received the booster dose. (**Figure 2H**). Significantly higher antibody levels were also observed at 6mopp among those initially vaccinated at T3 and then received a booster (*p*=0.0174). Antibody levels in women who were vaccinated post-partum increased post booster dose at 9mopp (*p*=0.0622) and 12mopp (*p*=0.0536). Breastmilk antibody titers significantly positively correlated with plasma antibody titers (**Figure 2I**) at 6 weeks (r=0.1346 *p*=0.0039) and again after (55.27 days) receiving the booster at 12mopp (r=0.3925 *p=*0.0041).

**Figure 2:**
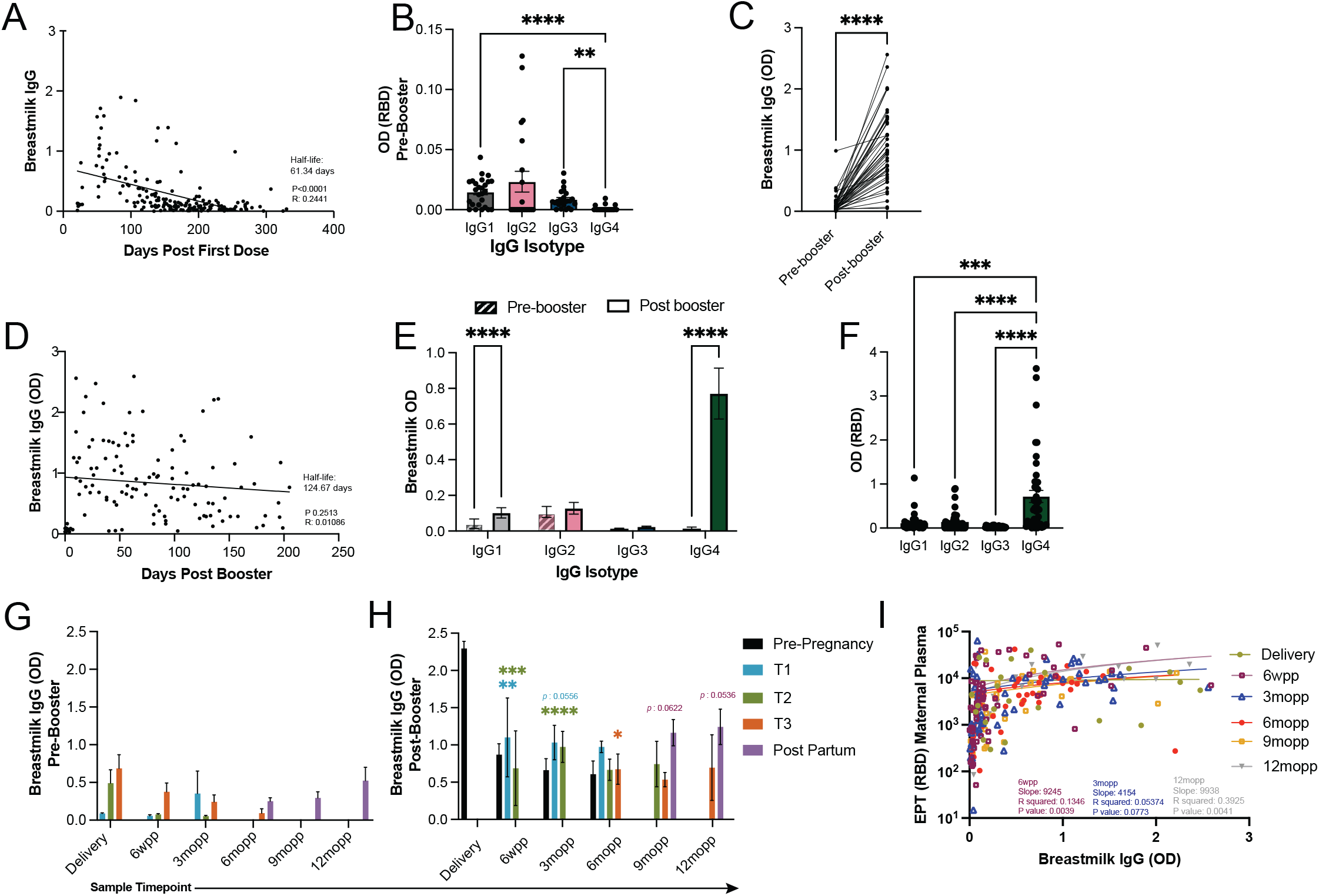
SARS-CoV-2-specific IgG antibody titers in breastmilk correlate with those in maternal circulation. **(A)** RBD-specific IgG levels in breastmilk after the first and second vaccine doses (n=179). **(B)** IgG isotype levels in breastmilk 174.33 ± 10.08 days after the first and second vaccine doses, (IgG1, n=26; IgG2, n=22; IgG3, n=24; IgG4, n=22). **(C)** Breastmilk IgG levels 37.84 ± 3.80 days prior to and 55.32 ± 5.30 days after booster (n=45 pairs). **(D)** RBD-specific IgG antibodies in breastmilk after maternal booster vaccination (n=123). **(E)** Levels of RBD specific IgG isotypes in breastmilk 57.50 ± 8.17 days before (n=28) and 117.23 ± 11.32 days after the booster dose (n=44). **(F)** IgG isotype detection of SARS-CoV-2 RBD specific antibodies in breastmilk 115.10 ± 11.46 days after booster dose (n=43). **(G-H)** Average IgG titers post-partum in women who had not received the booster dose **(G)** (Prepreg, n=0; T1, n=15; T2, n=52; T3, n=51; Postpartum, n=39) and **(H)** who had been boosted (Prepreg, n=26; T1, n=14; T2, n=38; T3, n=21; Postpartum, n=12) classified by trimester of initial vaccination. * indicates a significant difference between the antibody levels at that timepoint when comparing pre- and post-booster (panel G vs H). Data are median values ± SEM. **(I)** Correlation between RBD-specific IgG levels in breastmilk and maternal plasma post-partum (Delivery, n=24; 6 weeks (6wpp), n=60; 3 months (3mopp), n=59; 6 months (6mopp), n=48; 9 months (9mopp), n=32 and 12 months (12m), n=19). ∗ *p* < 0.03, ∗∗ *p* < 0.002, ∗∗∗ *p* < 0.0002, ∗∗∗∗ *p*<0.0001.

### Maternal IgG antibodies are passively transferred and are detected in cord blood plasma

At delivery, RBD-specific IgG antibodies were detected in umbilical cord blood (UCB) plasma albeit at significantly lower levels than in maternal circulation at delivery (*p=*0.0012) or peak maternal IgG titers (**Figure 3A**,**B**). Significant difference was most evident when mothers received their initial SARS-CoV-2 vaccination in the second (*p =* 0.0494) and third trimester (*p =* 0.0012) **(Figure 3C)**. RBD-specific IgG2 antibody titers were lowest in cord blood **(Figure 3D)**. Interestingly, there was no correlation between UCB RBD-specific IgG titers and maternal titers at delivery **(Figure 3E)**, peak maternal IgG levels pre-delivery (**Figure 3F**), or time since maternal first vaccination (**Figure 3G**).

**Figure 3:**
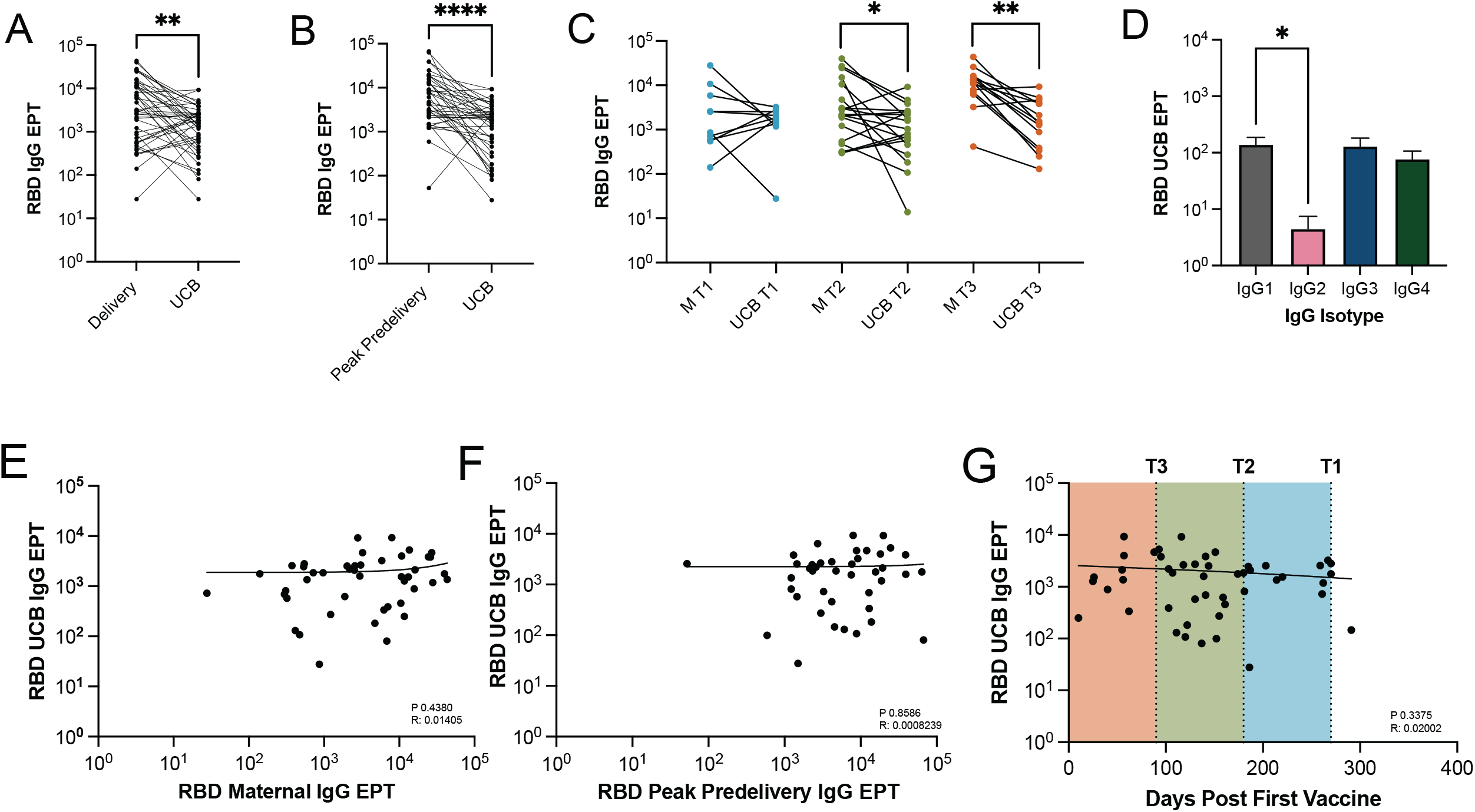
Maternal IgG antibodies generated in response to vaccination are detected in umbilical cord plasma. **(A)** RBD-specific IgG titers in maternal circulation and umbilical cord plasma at delivery (n=45 pairs). **(B)** Correlation of peak levels of RBD-specific IgG antibodies in maternal circulation pre-delivery and UCB (n=41). **(C)** Comparison of antibody levels in UCB and maternal circulation (M) at delivery, by trimester of initial maternal vaccination (T1=10, T2=19, T3=14). **(D)** IgG isotype analysis of UCB (n=14 pairs). **(E)** Correlation between UCB and maternal RBD-specific IgG titers at delivery (n=45). **(F)** Correlation between UCB and peak RBD specific IgG in maternal circulation before delivery (n=41). **(G)** RBD-specific IgG titers in UCB relative to days since maternal first vaccine dose (n=48). ∗ *p* < 0.03, ∗∗ *p* < 0.002, ∗∗∗ *p* < 0.0002, ∗∗∗∗ *p*<0.0001.

### Maternal IgG antibodies are present in newborn circulation

UCB is often used as surrogate for newborn blood,^38, 39^ however, there may be key differences in antibody transfer into cord blood and fetal circulation. Therefore, we next assessed RBD-specific IgG titers in newborn blood. IgG antibody titers were comparable in paired UCB and newborn plasma samples (**Figure 4A**) and correlated with each other (r=0.1943 *p=0*.0056) (**Figure 4B**). Moreover, as described for UCB plasma, RBD-specific IgG titers in newborn plasma were lower than those observed in maternal circulation at delivery (*p=*0.0200) as well as compared to peak maternal levels pre-delivery (*p<*0.0001) (**Figure 4C, Supp. 1A**), especially for mothers who received their initial vaccination series during the third trimester (*p*=0.0200) **(Figure 4D)**. However, in contrast to UCB, a significant positive correlation was observed between newborn plasma and paired maternal plasma RBD-specific IgG titers at delivery (r=0.3782 *p=*<0.0001) (**Figure 4E**), but not peak maternal levels pre-delivery (**Supp. 1B**). As described for UCB, titers of RBD-specific IgG2 titers were lowest in newborn plasma (**Figure 4F)**. Unlike UCB, newborn antibody titers were significantly inversely correlated with the time since initial maternal vaccination (r=0.3130 *p=*0.0002) (**Figure 4G**) with lower newborn IgG antibody titers for infants born to mothers vaccinated in early pregnancy.

**Figure 4:**
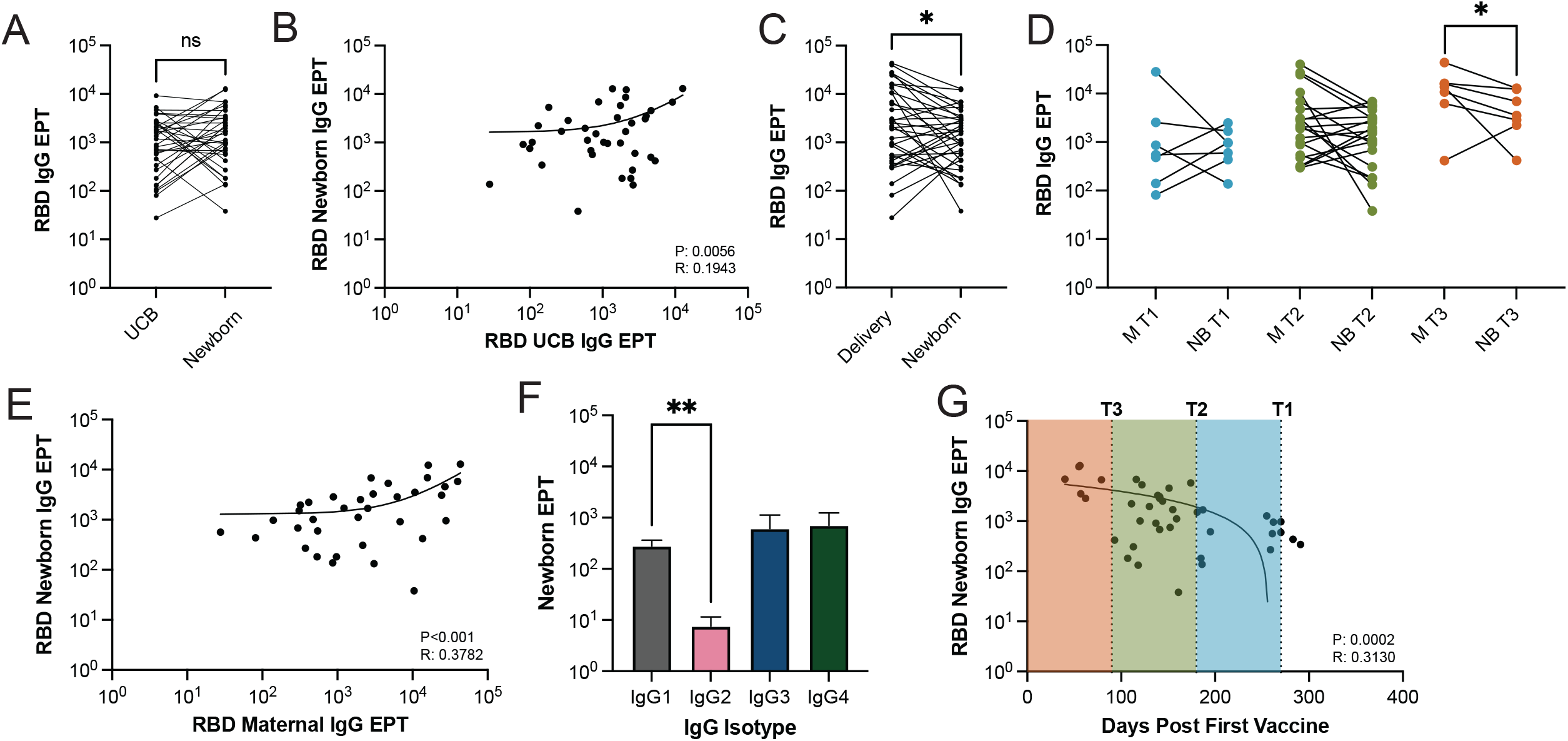
Passively transferred antibodies are detected in newborn circulation. **(A)** Comparison (n=38 pairs) and **(B)** correlation (n=38) between UCB and newborn blood RBD-specific antibody titers. (**C)** Overall comparison between maternal RBD-specific IgG antibodies at delivery and newborn RBD-specific IgG titers, independent of trimester of initial vaccination, (n=35) and **(D)** RBD-specific IgG titers in maternal plasma (M) at delivery and newborn plasma stratified by trimester of initial maternal vaccination (T1, n=6; T2, n=20; T3, n=7). **(E)** Correlation (n=35) of RBD-specific IgG titers in newborn and maternal plasma at delivery. **(F)** IgG isotype analysis in newborn plasma (n=14 pairs). **(G)** RBD-specific IgG titers in newborn plasma relative to days post maternal vaccination (n=40). ∗ *p* < 0.03, ∗∗ *p* < 0.002, ∗∗∗ *p* < 0.0002, ∗∗∗∗ *p*<0.0001.

### Impact of fetal sex

Previous studies have indicated that male fetal sex is associated with lower maternal EPT and transplacental transfer of SARS-CoV-2 antibodies following SARS-CoV-2 infection,^40^ therefore we investigated the impact of fetal sex. No significant difference in peak RBD-specific IgG in maternal circulation pre-delivery **(Supp 1C)**, maternal antibody levels at delivery **(Supp 1D)**, UCB **(Supp 1E)**, or newborn plasma was observed based on fetal sex (**Supp 1F**).

### Comment

#### Principal Findings

It is well established that maternal vaccination during pregnancy is an effective method to protect neonates via passive transfer of maternal antibodies.^8, 13, 41^ Despite studies on the immunogenicity and efficacy of the SARS-CoV-2 vaccines in adult populations, vaccine hesitancy remains relatively high among pregnant women. ^24^ Our results confirm earlier conclusions that the initial two dose vaccination series during gestation resulted in appreciable RBD-specific IgG response in maternal circulation, UCB, and breastmilk. ^20, 27^ Importantly, longitudinal analysis of post-partum samples indicates that the booster dose is essential for producing higher and more durable antibody levels in both maternal circulation and breastmilk,^28, 29, 42^ and should be strongly encouraged for all pregnant people to increase neonatal passive immune protection against SARS-CoV-2.

#### Results in the Context of What is Known

Although SARS-CoV-2 RBD antibodies were present in UCB, their levels were significantly lower compared to maternal plasma. This observation is in line with previous studies of SARS-CoV-2 maternal infection ^31, 43^ and maternal vaccination,^27^ but differ from prior studies on Tdap vaccination ^44^ and another SARS-CoV-2 study that reported high antibody levels in UCB when compared to maternal.^32^ Furthermore, our data showed no correlation between maternal and UCB IgG titers against RBD. These data differ from a study that showed early third trimester vaccination resulted in the highest maternal antibody titer.^45^ A possible explanation for the discrepancy between our study and this earlier one may be differences in sample size (121 versus 1536). Moreover, considerable differences in the platforms used to measure antibody responses to SARS-CoV-2 mRNA vaccination ranging from traditional end-point ELISA to Luminex based antibody levels and OD measurements at one given dilution could be another explanation for the discrepancies between the results described herein and elsewhere. In addition, earlier studies reported a significantly higher level of IgG antibodies in arterial relative to venous cord blood.^46^ Thus, another possible explanation for the discrepancies between these studies and ours could stem from a variability in UCB sample collection.

Similar to our observations with cord blood plasma, IgG titers were also lower in newborn plasma compared to those in maternal circulation. In contrast to the data obtained using cord blood, we do see a significant negative correlation between the days post first vaccination and RBD IgG in newborn plasma, suggesting higher newborn EPT are associated with maternal vaccination in T3. The increased antibody presence in newborn circulation following vaccination during T3 agrees with the rationale driving current Tdap vaccination recommendation during gestational T3.^13, 47^ Furthermore, IgG1 titers in newborn circulation were higher than those observed in maternal circulation confirming the greatest transplacental transfer ratio of IgG1.^46^ These data highlight the need to examine newborn blood samples when feasible.

The booster resulted in a striking increase in antibody levels in breastmilk, independent of the trimester when the initial vaccine series was administered in line with results from previous studies.^29, 45^ We report a significant increase across all isotypes with IgG1 and IgG4 becoming dominant after booster vaccination. Plasma observations are similar to those reported in recent studies.^42^ Since B-cells undergo the class switching pattern of IgM>IgG3>IgG1>IgA>IgG2>IgG4, ^48^ the IgG4 dominance we observed suggest enhanced maternal B-cell class switching. Our study results align with a study that showed an increase in IgG4 with pregnancy and higher IgG4 response in the pregnant population when compared to non-pregnant individuals.^49^

#### Strengths and Limitations

Our study leverages paired longitudinal samples from subjects ranging from their initial SARS-CoV-2 vaccine through the administration of the booster dose. Furthermore, the collection of newborn blood at delivery allowed us to directly evaluate passive transfer into fetal circulation and draw comparisons to cord blood, a commonly used surrogate. However, our study is not without limitations, including its sole focus on RBD-specific IgG binding antibody responses targeting the sequence from the USA-WA1/2020 isolate, as well as a lack of functional assays to assess vaccine-induced virus neutralization and antibody Fc-dependent functions.

#### Clinical Implications and Conclusion

Taken together, our results show that SARS-CoV-2-specific maternal antibodies generated via vaccination are passively transferred *in utero* and after birth via breastfeeding but wane within 6 months after first vaccination dose. Furthermore, our longitudinal maternal data indicate that breastmilk antibody levels are dramatically increased by the booster dose. Therefore, the best protection against SARS-CoV-2 mothers can give to their offspring is to receive the 3-dose vaccination series including the booster dose at any point during pregnancy and prior to delivery to allow for placental antibody transfer, and to subsequently breastfeed their children for at least 6 months, at which point their infants are eligible for vaccination as the CDC recently authorized SARS-CoV-2 vaccination of children starting at 6-months of age. ^50^ Continued breastfeeding throughout the first year of life is encouraged as SARS-CoV-2-specific maternal antibody levels persist in breastmilk following booster dosing for at least 12 mopp.

## FIGURE LEGENDS

**Supplemental Figure 1:**
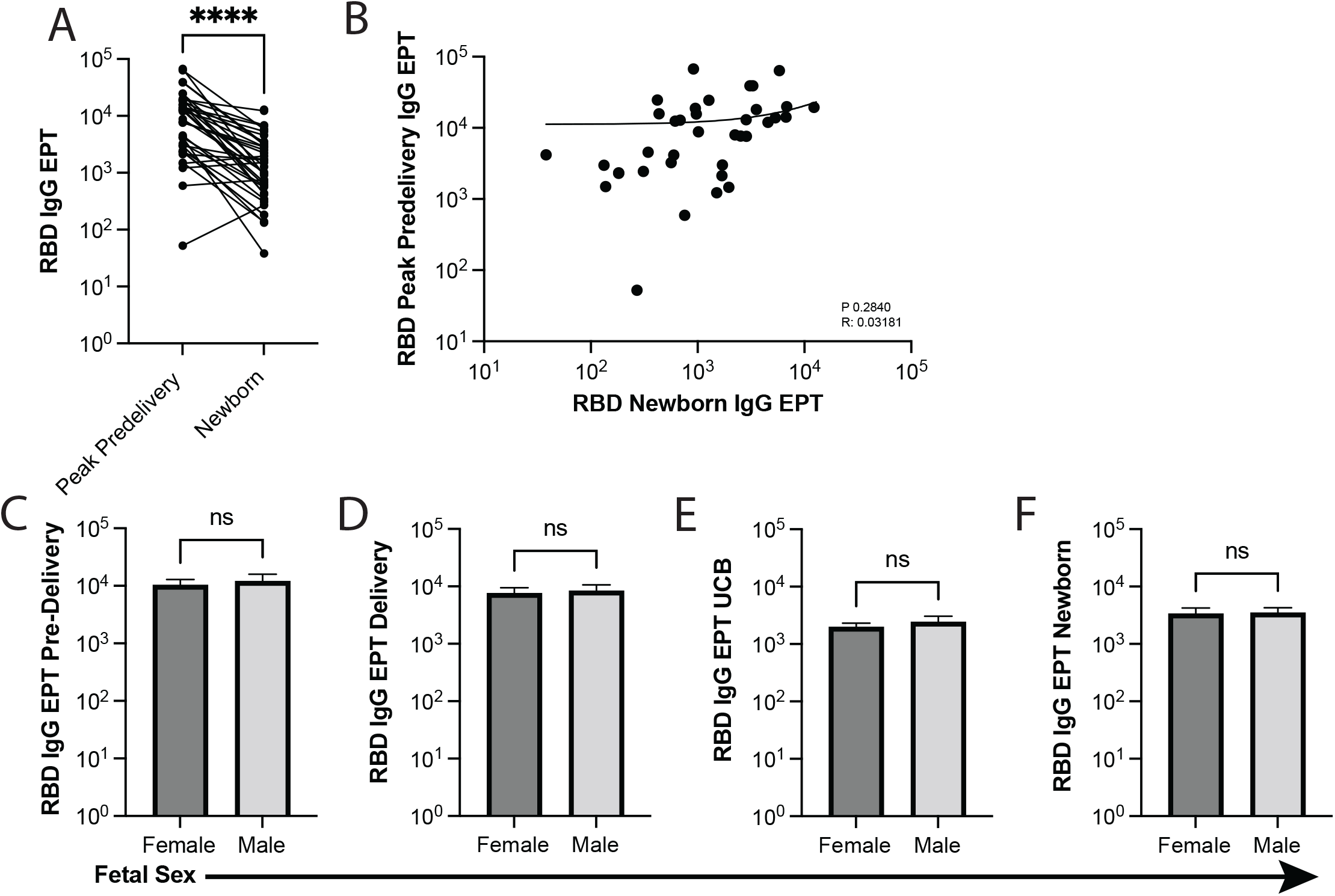
Fetal sex does not influence maternal antibody response to the SARS-CoV-2 vaccine or the passive transfer of SARS-CoV-2 vaccine-elicited antibodies. **(A)** Comparison (n=38) and **(B)** correlation (n=38) between peak RBD-specific IgG titers in maternal circulation pre-delivery and newborn plasma. **(C)** Peak maternal antibody titers pre-delivery (Female, n= 28; Male, n=19). **(D)** Antibody EPT of maternal plasma delivery (Female, n=34; Male, n=25). **(E)** RBD-specific IgG titers in UCB (Female, n=31; Male, n=23), and **(F)** RBD-specific IgG titers in newborn plasma (Female, n=24; Male, n=23) by fetal sex. Data in bar graphs are mean ± SEM. **(G-J)** Comparison of IgG isotype antibody titers in newborn blood, UCB, and maternal plasma at delivery. ∗ *p* < 0.03, ∗∗ *p* < 0.002, ∗∗∗ *p* < 0.0002, ∗∗∗∗ *p*<0.0001.

